# Contribution of floral transmission to the assembly and health impact of bacterial communities in watermelon seeds

**DOI:** 10.1101/2025.02.12.637993

**Authors:** Gillian E. Bergmann, Kacie Lui, Carina Lopez, Alexander Velasco, Coralie Marais, Matthieu Barret, Marie Simonin, Rachel L. Vannette, Johan H.J. Leveau

## Abstract

Flower-sourced assembly of seed microbiota remains an understudied ecological process. Here, we investigated the floral transmission pathway for bacterial acquisition by developing seeds of watermelon (*Citrullus lanatus*). Comparison of stigma- and seed-associated bacterial communities from field-grown *C. lanatus* revealed significant overlap: up to 40% of the bacterial diversity that was detected in seed was also found on stigmas. In a field pollinator exclusion experiment, honeybee visitation to flower stigmas had no significant effect on bacterial community composition in seeds. Among a collection of bacterial isolates from stigmas and seeds in the field, more than half (57%) were able to transmit to seeds after inoculation onto stigmas under laboratory conditions. Interestingly, for most bacterial strains, fruit set rates increased after floral inoculation, and in some cases even in the absence of transmission to the seed. We also found that bacterial isolates from watermelon stigmas and seeds had variable (i.e. positive or negative) effects on seed germination and seedling emergence. Our findings highlight the contribution of floral transmission to seed microbiota assembly and its consequences for plant fitness.

## Introduction

Bacterial communities associated with seeds are acquired either from the parent plant (Sulesky-Grieb *et al*., 2024) or from the ambient environment (Yao et al. 2024). These seed-associated bacteria may be vertically transmitted to developing seedlings during emergence (Abdelfattah *et al*., 2021, 2022; Johnston-Monje *et al*., 2021). Because seed-borne bacteria can modulate seed germination (Tang *et al*., 2023) and seedling tolerance to pathogens (Pal *et al*., 2022; Luo *et al*., 2024), the outcomes of seed bacterial community assembly can have major consequences for plant fitness. For this reason, there is much interest in predicting and manipulating seed bacterial communities for improving crop establishment sustainability, and for ecological conservation and restoration (Busby *et al*., 2017, 2022; Singh *et al*., 2023). However, seed bacterial community assembly is a product of many processes that interact in space and time, resulting in communities that can vary dramatically and erratically between seeds and across biological and ecological scales (Bintarti *et al*., 2022; Chesneau *et al*., 2022; Kim *et al*., 2023; Yao *et al*., 2024). We posit that a clearer understanding of the early stages of plant microbiota assembly will benefit from identifying the sources and pathways by which seeds acquire bacteria and from knowing the impacts of these bacteria on seed fitness (Bergmann & Leveau, 2022).

Seeds acquire bacterial communities through various pathways (Hodgson *et al*., 2014). One of these is the floral transmission pathway, where bacteria from stigmas and other flower tissues disperse through the style to developing seeds (Maude, 1996). Such early colonization may affect the ability of other bacteria to colonize seeds later in development through priority effects (Debray *et al*., 2022). Previous studies have demonstrated that bacterial and fungal pathogens (Ngugi & Scherm, 2004; Lessl *et al*., 2007; Dutta *et al*., 2014; Darrasse *et al*., 2018) as well as beneficial bacteria (Mitter *et al*., 2017) can colonize seeds from flowers, which has considerable consequences for seed viability and pathogen spread during seed-to-seedling transmission. This suggests that floral transmission indirectly affects seedling fitness. To date, very few published studies have explored the overlap of bacterial communities between flowers and seeds, so the range of bacterial taxa that may colonize seeds via floral transmission is largely unknown. Chesneau et al. (2020) looked at two plant species, *Phaseolus vulgaris* and *Raphanus sativus*, and found that 10-30% of seed-associated species were also present in the microbiota of whole flowers. Further comparison of these communities, through complementary field and lab studies, will give broader insights into the contribution of floral transmission in seed bacterial community assembly and the ecological processes that might be involved.

If stigmas are a main source for seed microbes, then insect pollination could well affect microbial assembly in seeds. Bees are known to vector bacteria between flowers, which affects the composition of nectar communities (Vannette *et al*., 2012; Aizenberg-Gershtein *et al*., 2013; Ambika Manirajan *et al*., 2016; Rebolleda-Gómez & Ashman, 2019) as well as floral attractiveness to pollinators (Herrera *et al*., 2008; Vannette *et al*., 2012; Rering *et al*., 2018). It is conceivable that some of the bacteria deposited on stigmas by bees could also transmit into developing seeds. In one study, pollination was shown to introduce bee-associated taxa to seeds of *Brassica napus* and alter seed community composition (Prado *et al*., 2020). However, these experiments were conducted in the greenhouse with open bee pollination and a self-pollinating plant species. As such, more work is needed to account for the role of pollinator visitation in the acquisition of seed microbes under field conditions with a plant species that cannot self-pollinate.

Watermelon (*Citrullus lanatus*) is a tropical fruit crop produced worldwide, including the Central Valley of California (USDA/NASS, 2023). The plant has dimorphic flowers, with heavy pollination requirements from insects for seeds to develop (Wijesinghe *et al*., 2020). It is also a host for the seed-borne pathogen *Acidovorax citrulli*, which causes bacterial fruit blotch across cucurbit species (Burdman & Walcott, 2012). This pathogen infects developing seeds from flowers (Walcott *et al*., 2003; Lessl *et al*., 2007), with the ability to infect internal seed tissues (Dutta *et al*., 2012a). Given the pollination biology and thorough study of pathogen transmission through stigmas, watermelon is an excellent model for testing the ecological importance of the floral pathway. Work has been done on the bacterial communities of *C. lanatus* seeds (Khalaf & Raizada, 2016, 2018), but this was limited to culture-based surveys. The overlap between stigma and seed bacterial communities was not considered in that study, and the role of floral transmission remains unknown.

In this study, we used a combination of field, greenhouse and lab studies to address the role of floral transmission in seed bacterial community assembly and seed fitness, using *C. lanatus* as our model of choice. We sought to answer the following questions: Q1) What is the overlap between stigma- and seed-associated bacterial community composition, Q2) Does bee visitation alter the composition of seed bacterial communities, Q3) Can we demonstrate the floral transmission of bacteria experimentally, and Q4) What is the impact of florally transmittable bacteria (i.e. bacteria that can successfully transmit to seeds from stigmas) on seed fitness? To address Q1, we collected floral stigma and seed samples from four commercial watermelon fields for comparison using 16S rRNA gene amplicon sequencing. We addressed Q2 with *gyrB* gene amplicon sequencing of seeds collected from a field pollinator exclusion experiment. Finally, we used a culture collection of bacteria isolated from *C. lanatus* stigmas and seeds in the field in inoculation experiments on stigmas (Q3) and seeds (Q4).

## Methods

### Description of the study system

*C. lanatus* is a vinelike plant, with multiple stems that creep close to the soil (Free, 1993) and flowers that continually develop until fruits begin to mature (Lu *et al*., 2003). Female flowers have three large stigmata that are radially symmetrical, with each stigma corresponding to a carpel in the ovary (Mann, 1943). The pollen grains on male flowers are large, sticky and rarely carried by wind (Porter, 1933), so stigmas can only be sufficiently pollinated by insects (Free, 1993).

### Field descriptions and sample collection

In the summer of 2022, we sampled stigmas and fruit from four commercial watermelon fields in California for our microbiota survey (Q1): two fields in Colusa County (i.e. Co1, Co2) and two in San Joaquin County (i.e. SJ1, SJ2). During sampling, we collected metadata on field location and fruit cultivar (Table S1). We collected samples from 10 random plants in a Z-pattern throughout each field. We harvested stigmas (*n*=10 per field) by cutting them from the flower and then placing them in Eppendorf tubes in the field using sterile tools. On each plant from which we collected stigma samples, we flagged several of the open female flowers. Seven days later, we returned to the field to harvest fruits that had developed from those flagged flowers by cutting them at the stem with sterile scissors and placing them in sterile Ziploc bags (*n*=10 fruit per field).

### Pollinator exclusion field experiment

To test if bee visitation affects the diversity and composition of seed bacterial communities, we conducted a field experiment in the summer of 2023. We ran this experiment in a commercial field in Sacramento County, California, during a week-long round of pollination by commercial honeybee hives. We created four 6×6-m plots at random locations in the field to account for environmental variation. At the onset of anthesis, we randomly bagged within each plot the male and female flowers of at least six plants in each plot. Twenty-four hours later, we randomly assigned all bagged and open female flowers to two groups: a hand pollination treatment, and a hand+bee pollination treatment. For the hand pollination group, we collected a bagged and open male flower, used it to hand-pollinate a bagged female flower by dabbing the anthers against the stigma until a uniform layer of pollen grains was deposited across all lobes. The female flower was then immediately re-bagged. For the hand+bee pollination group, we hand-pollinated as in the control group, and then left the flowers unbagged to allow bee visitation. We watched these flowers until each was visited by four honey bees (one of the visits was by a *Leucopis* species), and then immediately re-bagged them. The choice of four visits was based on previous observations that between two and five bee visits are required to successfully pollinate a watermelon flower (Campbell et al. 2018). During pollination, we collected four bees per plot for comparing potentially shared bacteria by trapping them in 50 mL falcon tubes while they sat on flowers. Bees that were brought back from the field were euthanized by incubation at -20°C for 24 hours. Seven days after applying the treatments, we harvested the fruits for library preparation using the same protocol as described above. A total of 24 fruits were collected: 12 fruits per treatment, with three fruits per treatment in each plot.

### Preparation of the metabarcoding libraries

To prepare stigma samples for 16S rRNA gene amplicon sequencing, we used a modified version of the protocols from Cui et al. (2020a, 2020b): we added 200 µL of sterile 0.5X Phosphate-Buffered Saline (PBS) to each stigma in each tube, vortexed for five minutes, discarded the stigmas and used the resulting suspensions for DNA extraction (see below). To process seeds, we surface-sterilized fruits in a laminar flow hood by submersion in 1% sodium hypochlorite with 0.001% Tween 20 for one minute, then three times in sterile water for one minute each. After drying the fruits for 10 minutes, we cut each fruit into sections and removed seeds from each section. We lyophilized the seeds for 24 hours in a FreeZone Benchtop Freeze Dryer (Labconco, Fort Scott, Kansas). Bees from the pollinator exclusion experiment were pooled, suspended in 1X PBS with 0.001% Tween 20, and macerated, as per the protocol from Prado et al. (2020).

We extracted DNA from stigmas using a PowerSoil kit, and from seeds and bees using a DNeasy Plant Pro kit (QIAGEN, Germantown MD). Different kits were used for different tissue samples to achieve proper homogenization and removal of plant inhibitors. To get sufficient microbial DNA from seeds, we pooled up to 100 mg of dried seed material (i.e. 50-100 seeds) per single fruit. As a negative extraction control, we used the same extraction protocol on sterile water. DNA yield was verified via PCR and gel electrophoresis. DNA extracts from the field survey samples were sent to NovoGene (Sacramento, CA) for 16S rRNA gene library preparation using primers 799F/1193R (Beckers *et al*., 2016) during stage 1 PCR amplification of the V5-V7 region, followed by 250-bp paired-end sequencing on the Novaseq 6000 platform. DNA extracts from the pollinator experiment samples were sent to the ANAN platform at INRAE-IRHS (Angers, Maine-et-Loire, France) for 250-bp paired end sequencing of the *gyrB* gene on the Illumina MiSeq platform, using gyrB_aF64/gyrB_aR353 (Barret *et al*., 2015) primers. One negative PCR control (i.e. sterile water as PCR template) was included in each sequencing run.

### Processing of Illumina metabarcode libraries

We are aware that low-biomass samples such as stigma washes and seeds can be prone to contamination during metabarcode library preparation (Karstens *et al*., 2019). As such, we included multiple negative controls (described above) and strict filtering protocols during bioinformatic processing (described below) to limit the presence of contamination in our datasets. We received the raw sequencing data for the field survey and the pollinator experiment in a demultiplexed format, trimmed off the primer sequences using Cutadapt v2.7 (Martin, 2011) and processed the resulting sequences using the DADA2 workflow (Callahan *et al*., 2016). Briefly, we filtered out low quality reads using a max expected error (maxEE) threshold of 2, denoised sequence composition in the samples and dereplicated duplicate sequences. We then assigned taxonomy to the resulting amplicon sequence variants (ASVs) with IDTaxa (Murali *et al*., 2018) using Silva release 138 (Pruesse *et al*., 2007) to identify 16S rRNA sequences, and train_set_gyrB_v5 to identify *gyrB* sequences. We then filtered out ASVs not identified as bacteria, along with ASVs identified as the *gyrB* paralog *parE.* After this filter, we compiled the sequence composition table, taxonomy, sequences and sample metadata for each dataset into phyloseq objects (McMurdie & Holmes, 2013) for additional filtering and downstream analyses.

Using these phyloseq objects, we filtered out putative contaminants using the prevalence method in Decontam v.1.24.0 (Davis *et al*., 2018) with a probability threshold of 0.2. We then removed samples with sequencing depths below 500 reads. Finally, we generated a subset phyloseq object with singletons removed and converted read counts to relative abundances in both objects. The outputs of this processing workflow were six phyloseq objects: two for the field survey and four for the pollinator experiment. The 16S rRNA dataset for the field survey contained 2,623,671 total reads, with an average sequencing depth of 34,522 reads per sample, representing a total of 1301 ASVs before filtering out singletons and 1259 ASVs after filtering, distributed across 30 stigmas samples and 32 seed samples. The *gyrB* dataset for the pollinator experiment had 292,560 total reads, with an average sequencing depth of 9,143 reads per sample. There were 2,117 ASVs prior to filtering out singletons and 546 ASVs after filtering, distributed across 4 bee samples and 18 seed samples.

### Analysis of field survey and pollinator experiment datasets

All analyses on the sequence-based field survey and pollinator exclusion experiment were done in R 4.4.0 (R Core Team, 2016). We first compared the alpha diversity in stigma and seed communities within and across fields from the 16S rRNA field survey (**Q1**). We used ANOVA and *post-hoc* pairwise comparisons for both observed ASV richness and Faith’s phylogenetic diversity (Faith’s PD) with tissue type as the predictor. We calculated Faith’s PD using a phylogenetic tree generated with phangorn v.2.11.1 (Schliep *et al*., 2023). We then compared community composition between stigmas and seeds using Non-Metric Dimensional Scaling (NMDS) ordination and Permutational Analysis of Variance (PerMANOVA) in vegan v.2.6.6.1 (Okansen *et al*., 2019). These analyses were based on a weighted Unifrac distance matrix with tissue type and field as predictors. We also compared communities between fields for each tissue type separately using NMDS ordination and PerMANOVA with field as the predictor. Finally, we calculated the Jaccard beta-diversity between individual pairs of stigmas and seeds and partitioned this diversity into its nestedness and turnover components with betapart v.1.6 (Baselga & Orme, 2012).

Next, we identified taxa shared generally across communities using UpSetR v.1.4.0 and ComplexUpset v.1.3.3 (Conway *et al*., 2017; Krassowski, 2020) with tissue type as the intersection variable. In addition to calculating the overlap across all stigma and seed samples, we also compared stigmas and seeds that were sampled from the same plant to measure the variation in the number of ASVs shared and their cumulative relative abundance in each tissue type. We subset the ASVs in our phyloseq object based on which tissues they were found in and visualized the dominant genera that were shared or unique to seeds with microbiome v.1.26.0 (Lahti & Shetty, 2017). Finally, we identified shared genera that were enriched or depleted in seeds compared to stigmas using differential abundance analysis with ANCOM-BC2 v.2.6.0 (Lin & Peddada, 2020).

For the pollinator experiment, we used similar methods to test for differences in alpha diversity and community composition of seed bacteria between flowers that were or were not allowed bee visitation (**Q2**). We also identified overlap in taxa between bees and seeds within each treatment group. We first compared Faith’s PD alpha diversity in seeds between treatments using a Student’s t-test. We compared seed communities between treatments with NMDS ordination and PerMANOVA of weighted Unifrac distances with treatment group as the predictor. Finally, we generated two upset plots: one comparing taxa in bees and seeds from the hand-pollination group, and one comparing bees and seeds from the hand+bee pollination group. We also examined which taxa were shared between tissues within each treatment group, as well as the relative abundances of those shared taxa.

### Isolating stigma- and seed-associated bacteria from the field for inoculation experiments

In addition to our sequence-based field surveys, we generated a culture collection of stigma- and seed-associated bacteria for use in floral transmission experiments. For this, we collected stigmas and fruits from two commercial watermelon fields (Tule, Poundstone) in Colusa County, California in the summer and fall of 2021. Different cultivars were grown in each field: Charleston Gray in Tule, AU Producer in Poundstone. Using the same methods as described above, we collected ten stigmas per field, as well as fruits (five from Tule, nine from Poundstone) 53 days after stigma sampling. Stigmas and fruits were stored at 4°C until processing. We processed samples and isolated bacteria on 0.1X Tryptic Soy Agar (TSA) amended with 0.01 mg of benomyl per mL to minimize fungal growth. A detailed protocol for the isolation process is described in Supplementary methods 1.

From 72 morphologically distinct bacterial strains in our collection (33 from stigma, 39 from seed), we extracted DNA using REDExtract-n-Amp (MilliporeSigma, Burlington, MA) and NucleoSpin Microbial DNA (Macherey-Nagel, Allentown, PA) kits. We then ran PCR reactions containing 10 µL of GoTaq Green, 1 µL of 27F/1492R primers (Heuer *et al*., 1997), 4 µL DNA template and 9 µL water for a total reaction volume of 25 µL, under the following conditions: 95 °C for 4 minutes, 25 cycles of 95 °C for 30 s, 55 °C for 30 s and 72 °C for 60 s, followed by a final elongation at 72 °C for 10 s. We checked for amplification with gel electrophoresis, and amplicons of the correct size (1.5 kb) were purified with a Qiaquick PCR purification kit (QIAGEN, Germantown MD) and sent to the UC Davis and UC Berkeley DNA Sequencing labs for Sanger sequencing with the 27F primer. The resulting sequences were trimmed in Benchling (https://www.benchling.com/) and assigned taxonomy to the species level based on querying against type strains in NCBI’s Genbank (https://blast.ncbi.nlm.nih.gov/Blast.cgi).

### Stigma inoculation experiment

To validate the ability of stigma- and seed-associated bacteria to transmit from stigmas to seeds (**Q3**), we selected five strains from our culture collection (Table 1) for inoculation experiments. We used sterile 1X PBS as a negative control and two strains of *Acidovorax citrulli* (AAC00-1 and M6, isolated from seed and fruit respectively; Walcott *et al*., 2000; Burdman *et al*., 2005) as positive controls. We generated rifampicin-resistant variants of each strain in order to selectively culture them from seeds. We did this by making overnight liquid cultures in 0.1X TSB+0.01mg/mL benomyl or King’s B (KB; King *et al*., 1954) from a single colony of each strain and spread plating 100 µL of culture onto plates of KB amended with 100 *μ*g/mL of rifampicin. We incubated the KB+rif plates for 72 hours and re-streaked one of the appearing colonies for each strain onto another plate of KB+rif to verify resistance. We grew watermelon (cv. Sugar Baby) plants from seed (Burpee Seed Co, Warminster Township, PA) in a CMP6060 growth chamber (CONVIRON, North Dakota, USA) at 28°C and 80% relative humidity (RH) with 12h/12h light/dark cycles. We chose to use Sugar Baby for this experiment because it has been used in previous studies on watermelon seed pathogens (Johnson *et al*., 2011; Johnson & Walcott, 2013) and produces smaller i.e. more manageable plants.

When the plants began flowering, we prepared liquid cultures of each strain in KB+rif and grew them overnight in a rotary shaker at 270 rpm and 28°C. After incubation, we centrifuged for 5 minutes at 6,000 x *g*, poured off most of the supernatant, resuspended the cell pellet in PBS with 0.001% Tween 20. We quantified cell concentrations with a GENESYS™ 40/50 Vis/UV-Vis spectrophotometer (Thermo Fisher, Waltham, MA) and diluted each inoculum solution to 10^8^ cells/mL. We randomly assigned an inoculation treatment to each hand-pollinated female flower, and applied the inoculum by pipetting 10 µL of a cell suspension across the stigma surface. We continued to inoculate stigmas until we had acquired 4 developing fruits per treatment for analysis.

Seven days post-inoculation, developing fruits were collected and surface-sterilized for seed extraction as described above. To account for the radial symmetry of the stigmas and fruits, we vertically sliced each fruit down the midpoint between the stigma and the stem and then sliced each half in-between each carpel (Fig. 5A). We then removed and dried seeds as we did in the field studies. While the seeds dried, we prepared 96-well plates with 200 µL of KB+rif in each well, placed a single seed into each well, with 8 seeds sampled per fruit slice (i.e. 48 seeds per fruit, 2 fruits per plate), and sealed the plates with parafilm for incubation in a rotary shaker at 270 rpm and 28°C for five days. Turbidity in a well was recorded as positive for bacterial presence. With the number of positive seeds per fruit slice, we generated a heatmap of the average bacterial re-isolation frequency from seed per fruit slice for each strain in R 4.4.0 (R Core Team, 2016). We also compared the proportions of successfully developing fruits between inoculation strains using a chi-squared test and pairwise comparisons.

### Seedling viability experiment

To test the effect of bacteria on seedling fitness (**Q4**), we selected three isolates from our collection for which we had shown re-isolation from seeds after inoculation on stigmas in the previous experiment (see above), as well as both *Acidovorax citrulli* strains. As additional bacterial controls, we selected three strains from our culture collection, each one representing a genus that was not tested in the stigma inoculation experiment. We tested 8 strains in total. Finally, we included sterile, non-inoculated and non-sterile, non-inoculated seeds as controls.

For this experiment, we followed a protocol modified from Chesneau et al. (2022a) using watermelon (cv. Sugar Baby) seeds purchased from Burpee Seed Co (Warminster Township, PA). We inoculated seeds in batches of 25 each, with three replicate batches per treatment. We surface-sterilized seeds (except the non-sterile control batches) by washing them for 1 minute in 1% NaOCl with 0.001% Tween 20, followed by three washes in sterile deionized water for 1 minute each. After drying the seeds for 10 minutes, we soaked each seed batch for 30 minutes in a 10^7^ cells/mL cell suspension of each bacterial strain. We then dried seeds again under sterile conditions for 10 minutes. We transferred all seeds to culture tubes containing 5 mL of Murashige and Skoog plant nutrient medium (Murashige & Skoog, 1962) and incubated them in a Norlake Scientific plant incubator (Hudson, WI) at 28 °C, 80% RH and 12h/12h light/dark cycles.

To collect seedling fitness data, we recorded germination every 24 hours for seven days. If seedlings emerged, we incubated them for an additional seven days and then recorded their phenotypes as “normal” or “abnormal” based on rules from the International Seed Testing Association (https://www.seedtest.org/en/home.html). We compared the proportions of germinated seeds and normal seedlings between inoculation treatments in R 4.4.0 (R Core Team, 2016) with chi-squared tests and post-hoc pairwise comparisons.

## Results

### Stigma and seed bacterial communities overlap significantly (Q1)

Across the four commercial watermelon fields that were sampled, we found that bacterial species richness was higher on individual stigmas than in pooled seeds (t-test: t = 3.35, p = 0.002; Fig 1A), as was phylogenetic alpha diversity (t-test: t = 2.35, p = 0.02; Fig. S1). We also found that stigma and seed communities had significantly different compositions, but that tissue type only explained 6% of the variation between communities (Fig. 1B). This indicates that while stigma and seed communities host distinct taxa, a large subset is shared between tissues. When comparing communities within each tissue type, we found that source field affected the composition communities of both tissues (Fig. 1C). Field effects explained more of the variation in stigma microbial communities than in seed communities (Fig 1C).

**Figure 1.**
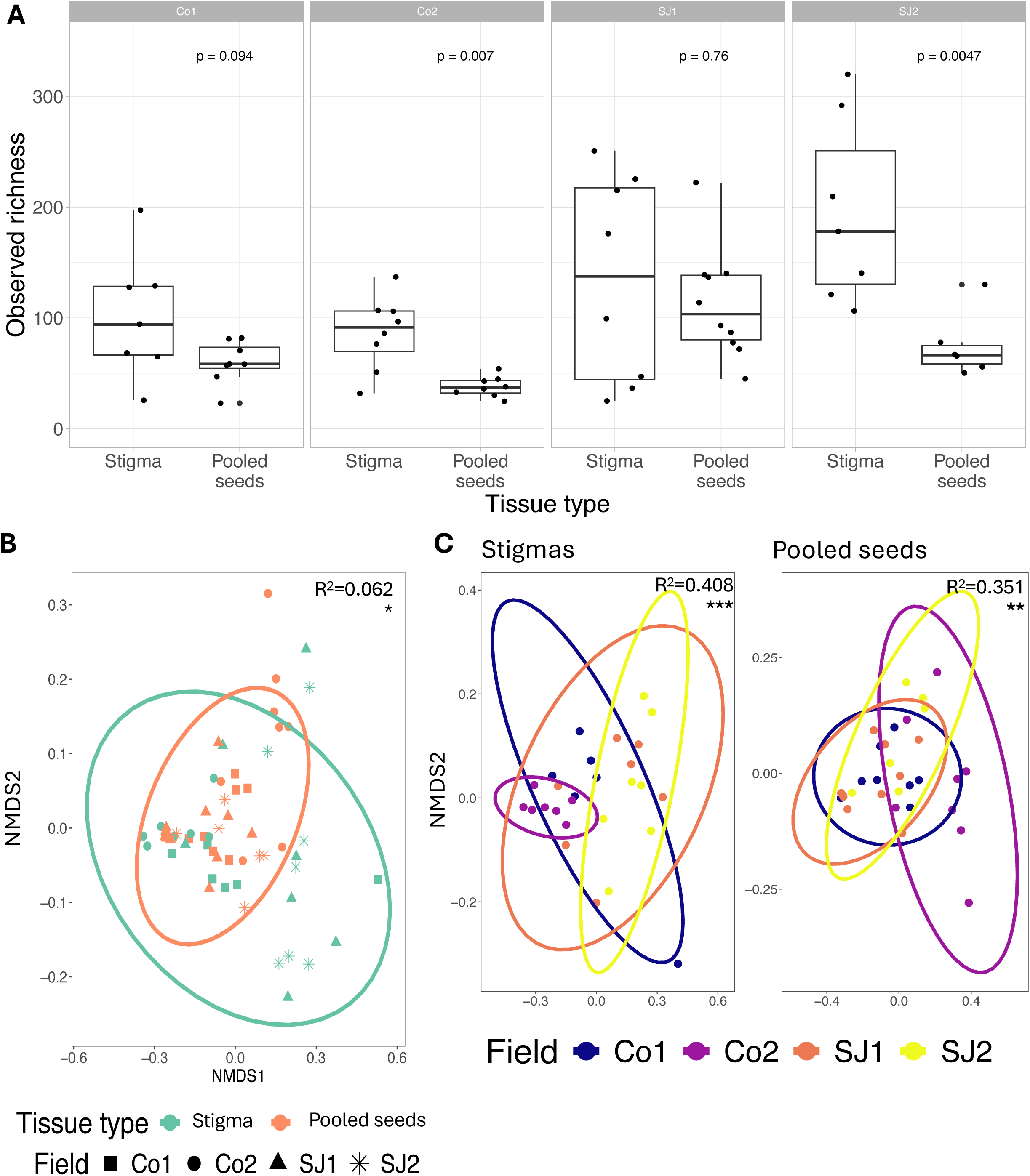
Seed communities are less diverse (A) and compositionally similar (B) to stigma communities. The boxplots show the variation of observed ASV richness (A) in stigma and seed communities in each sampled field. Each facet (Co1, Co2, SJ1, SJ2) represents a different field. The Non-metric dimensional scaling (NMDS) ordination plots of abundance-weighted Unifrac distances show the differences in community composition between tissues (B) and between fields within each tissue (C). These plots were generated from 16S rRNA amplicon sequencing data.

Stigmas and seeds shared 425 bacterial ASVs out of the 1259 observed (Fig. 2A). These shared ASVs represent 38.6% of all ASVs detected on stigmas and 73% of all ASVs detected in seeds. When we compared the bacteria found in seeds to those in individual stigmas from the same plant, we found that these shared bacteria make up 1.5-40% of the observed richness and 0.4-81.9% of the reads in seed communities (Fig. 2B-C). According to Jaccard index, turnover was the main component of community membership (Fig. S3A). However, this high turnover rate was mainly due to rare taxa (Fig. S3B).

**Figure 2.**
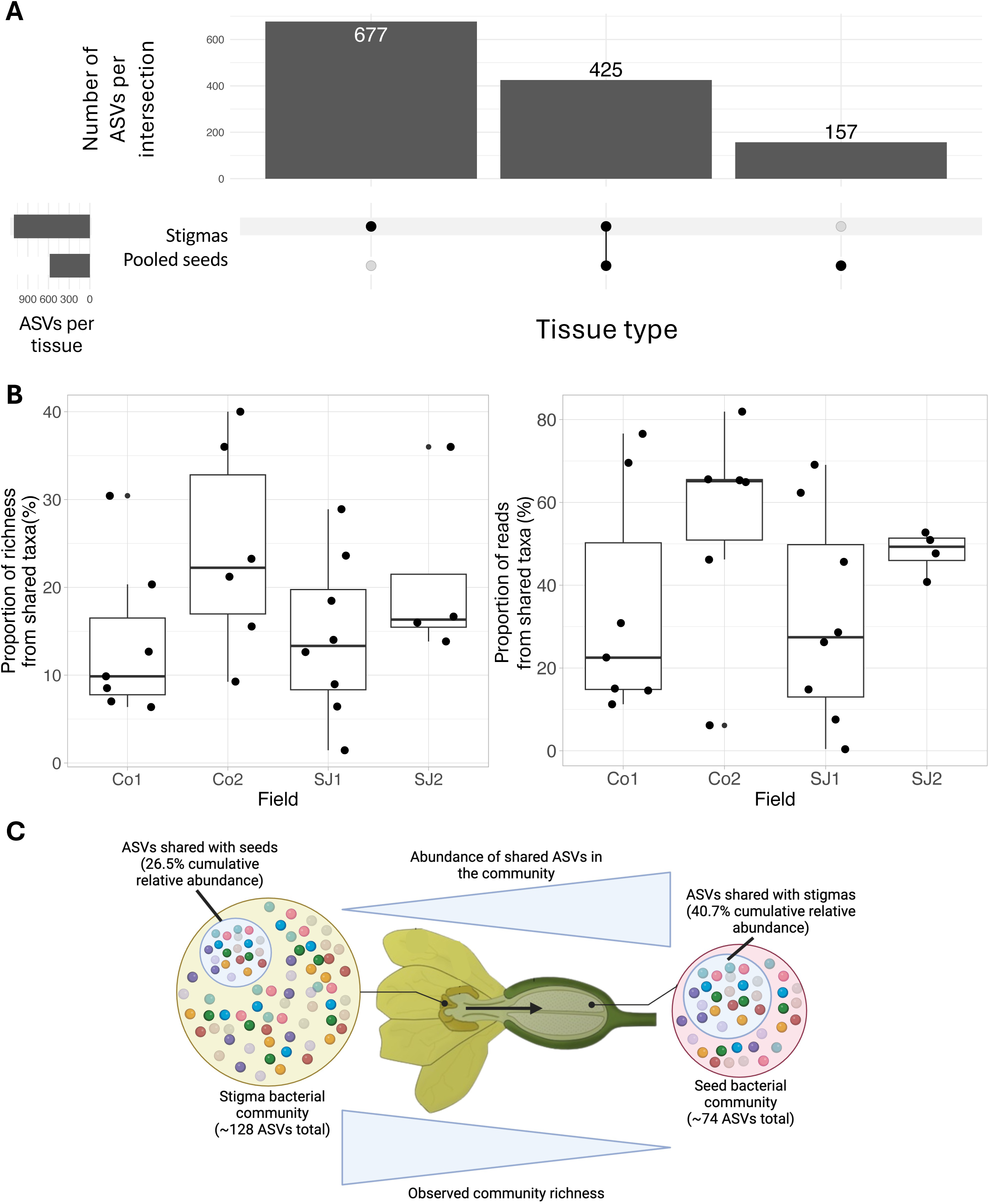
Shared taxa make up a large portion of stigma and seed communities. The UpSet plot of the number of ASVs detected in each tissue type (A). The richness proportions (B) were calculated by dividing the number of seed ASVs shared with stigmas by the total number of seed ASVs. The reads proportions (B) were calculated by summing the relative abundances of all shared ASVs for each seed sample. The overall trends of these richness and abundance data are summarized in the conceptual diagram (C). These plots were generated from 16S rRNA amplicon sequencing data.

When comparing the taxonomic composition of stigma and seed bacterial communities, we found that most shared ASVs belonged to the genera *Sphingomonas, Arsenophonus, Shigella* and *Acinetobacter* (Fig. 3A). *Apibacter, Weissella, Saccharibacter, Enterobacter* were unique to stigma communities, whereas *Skermanella, Faecalibaterium, Prevotella, Bacteroides* were unique to seed communities (Fig. S4). Through differential abundance analysis of the shared ASVs, we found that *Shigella* and *Paracidovorax* (formerly *Acidovorax*; Du *et al*., 2022) were enriched in seeds as compared to stigmas, while *Nocardioides, Bradyrhizobium, Arsenophonus* and *Ralstonia* were depleted (Fig. 3B). In total, we found that a large proportion of bacteria are shared between stigma and seed communities across multiple scales (field, plant).

**Figure 3.**
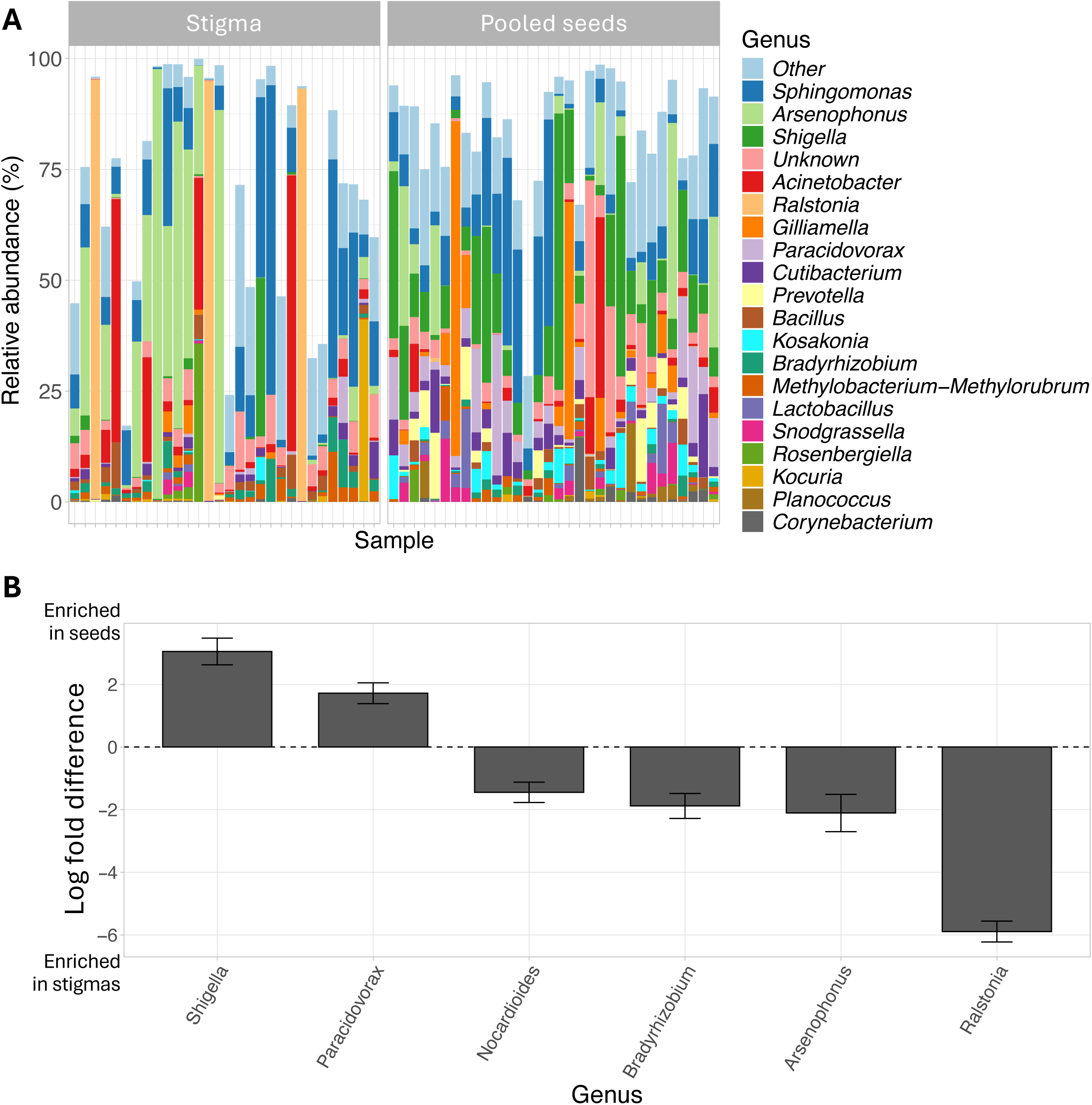
Genera shared between stigmas and seeds are generally abundant, with variation between tissues. We visualized the subset of bacterial communities that were shared between stigmas and seeds, aggregating to the top 20 most abundant genera in the subset (A). We then identified genera that are differentially abundant in seeds compared to stigmas (B). These comparisons were made using the subsets of taxa identified in the overlaps between all stigma and seed samples (i.e. not the pairwise comparisons). These plots were generated from 16S rRNA amplicon sequencing data.

### Bee visitation did not affect community composition of seed bacteria (Q2)

Given that *C. lanatus* is exclusively pollinated by bees and that some of the shared bacterial taxa observed in the field survey were bee specialists (e.g. *Gilliamella, Snodgrassella*), we tested if bee visitation impacted seed bacterial community composition in a separate field experiment. Specifically, we expected that seeds in a hand+bee pollination treatment would show more bee-associated ASVs and differ in diversity and composition compared to a hand-pollination-only treatment. When we calculated the overlap of ASVs between seeds and bees, we found that the same number of ASVs were shared across tissues in both treatments (Fig. 4A). When we compared alpha diversity between treatments, we found that bacterial diversity was slightly lower in the hand+bee-pollinated seeds, but this was not significant (Fig. 4B). Similarly, we found that the variations in community composition did not differ between treatments (Fig. 4C). The lack of significant seed bacterial communities indicates that bee visitation did not impact seed microbiota under our experimental conditions.

**Figure 4.**
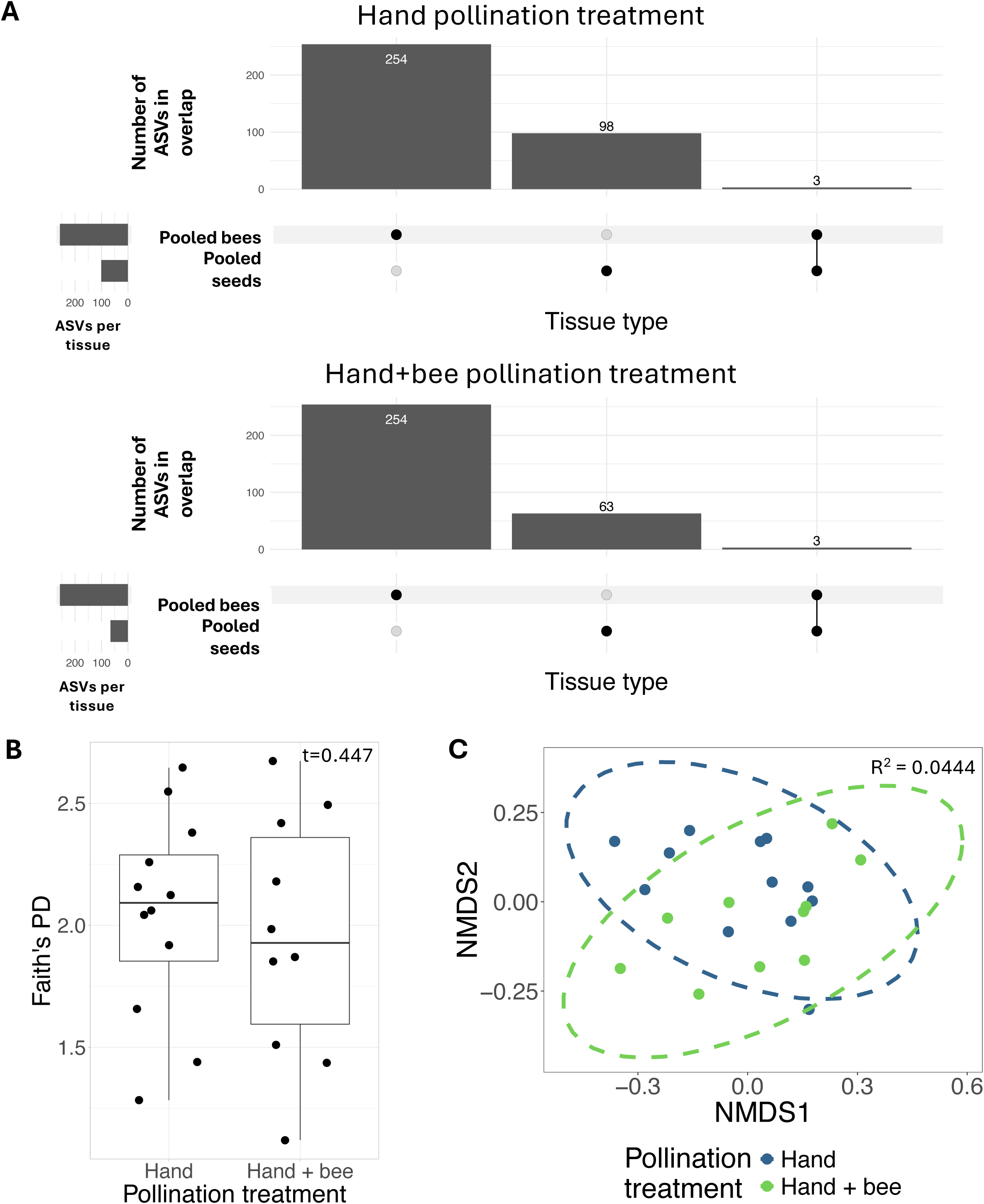
There was no significant variation in taxa overlap (A), diversity (B) and community composition (C) between pollination treatments. We compared Faith’s PD alpha diversity values between hand and insect pollinated seeds, as well as community composition using non-parametric dimensional scaling of abundance-weighted Unifrac distances. These plots were generated from *gyrB* amplicon sequencing data.

### Some but not all stigma- and seed-associated bacteria florally transmit to developing seeds (Q3)

To experimentally validate that bacteria can use the floral pathway to colonize the seed, we generated a collection of 54 bacteria isolated from watermelon stigmas and seeds. The majority of our strains were identified as *Bacillus,* which was the only genus for which representatives were isolated from both stigmas and seeds (Table 2). While all other genera were only cultured from one tissue or the other, all were present in both tissues in the sequence-based field survey (Table 1).

From this collection, we selected 5 strains: one that was collected from seed (*Bacillus* T.m10.b1) and four from stigmas (*Erwinia* P.s8.b2*, Paraburkholderia* T.s8.b1*, Rosenbergiella* P.s10.b1 and *Microbacterium* T.s9.b4). When each of these 5 strains, as well as two model strains of *Acidovorax citrulli (*AAC00-1 and M6, originally isolated from watermelon seed and fruit, respectively) were inoculated individually onto watermelon stigmas, only four strains could be re-isolated from seeds: AAC00-1, P.s8.b2, T.s8.b1 and P.s10.b1 (Fig. 5A). The rates of re-isolation varied by strain between fruits and between slices within each fruit (Fig. 5A), with the isolates of *Paraburkholderia* and *Rosenbergiella* consistently present across seeds from different fruit slices, whereas *Erwinia* and *A. citrulli* AAC00-1R were re-isolated at low frequencies from seeds closer to the stigma (Fig. 5A). These data validate that some bacteria are florally transmittable to seeds.

**Figure 5.**
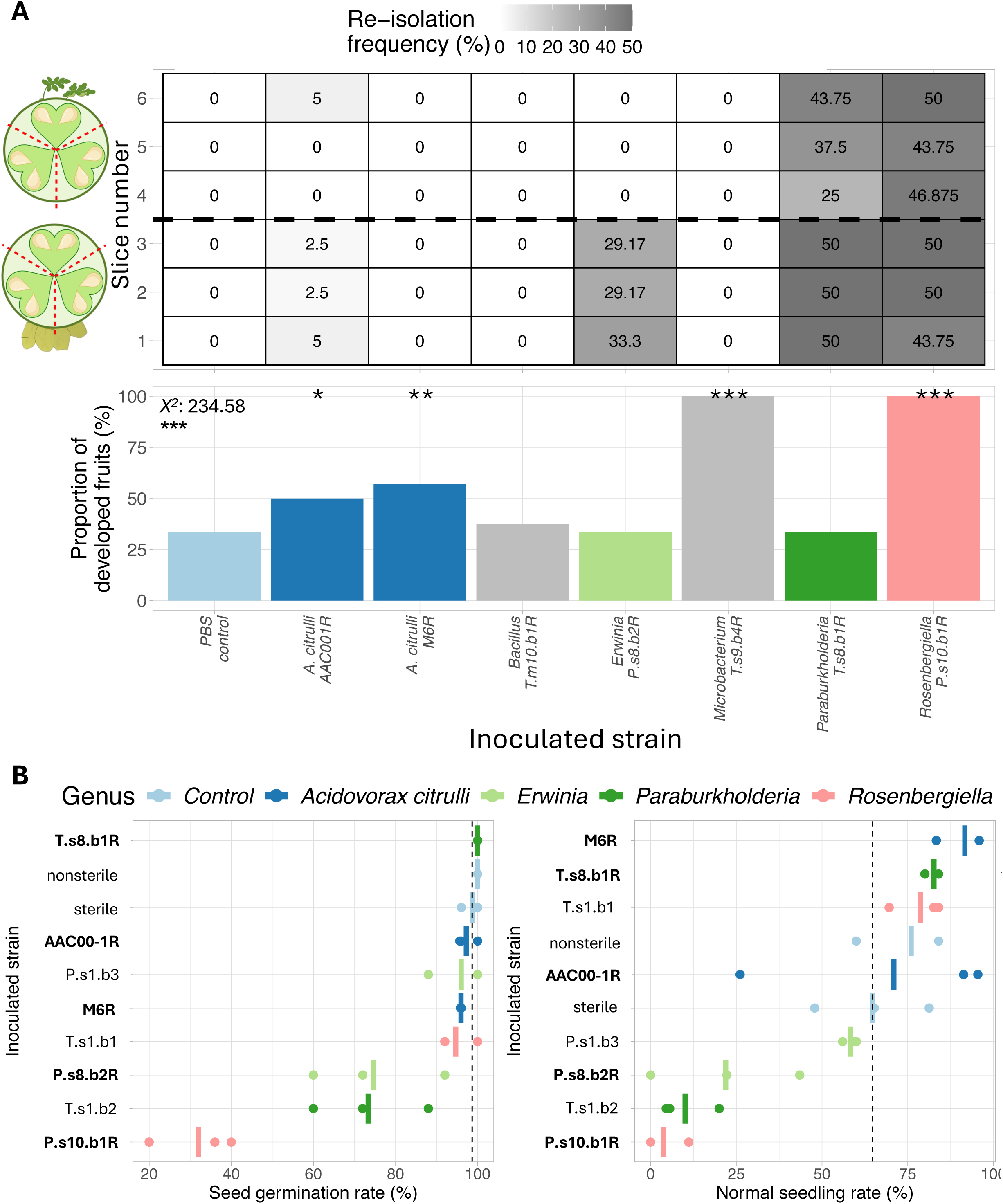
Bacterial strains had variable effects on floral transmission, fruit development and seed viability. The heatmap in (**A**) shows the average re-isolation frequency of bacterial strains from seeds collected from different fruit slices. The graphic next to the heatmap shows the spatial organization of seeds in the fruit slices, with the top image showing slices closer to the fruit stem and the bottom image showing fruit slices closer to the stigma. The barplot in (**A**) shows the proportion of fruits that successfully developed after stigma inoculation. Bars are color coded based on strain representation in **(B**) The dotplots in (**B**) show the seed germination and seedling development after inoculation with florally transmittable bacterial strains and their congeners. Strains named in bold were tested in the stigma inoculation experiment. The dotted lines represent the average value for each phenotype in the surface-sterilized control. Stars represent statistical significance based on residuals post-hoc pairwise comparisons.

### Florally transmittable bacteria impact fruit and seedling development (Q4)

In the stigma inoculation experiment, we observed that fruit set rates varied significantly between strains (Fig. 5A). Inoculation with *Rosenbergiella* improved fruit set by 67% compared to the PBS control, and inoculation with *A. citrulli* AAC00-1R improved fruit set by 17%. We also observed 24% and 67% increases in fruit set for stigmas inoculated with *A. citrulli* M6R and *Microbacterium sp.* T.s9.b4R respectively, although these bacterial strains were not detected in seeds from those inoculation experiments (Fig. 5A).

In a seedling viability test, we found that bacterial inoculation with seeds had significant effects on both seed germination and seedling development (Fig. 5B), with variation between strains. For example, compared to sterile controls, *A. citrulli* M6R increased normal seedling development by 27%, and *Erwinia sp.* P.s8.b2R reduced germination by 24% and normal seedling development by 43%. Interestingly, *Paraburkholderia sp.* T.s8.b1R improved normal seedling development while *Paraburkholderia sp.* T.s1.b2 reduced development. Similarly, *Rosenbergiella sp.* T.s1.b1 increased normal seedling development, while *Rosenbergiella sp.* P.s10.b1R greatly reduced both germination and development. These experiments show that florally transmittable bacteria and their conspecifics, many of which are assumed non-pathogenic, can have either positive or negative effects on fruit development and seed viability.

## Discussion

Through multiple experiments, we assessed the role of floral transmission in seed bacterial communities. In our field work, we found a significant overlap in taxa between stigma and seed communities, mainly representing genera that are commonly associated with the antho- and phyllosphere. In our subsequent lab studies, we found that some stigma- and seed-associated strains were florally transmittable. We also found that some of these transmittable strains had positive effects on fruit set (e.g. *Rosenbergiella*), germination and seedling development (e.g. *Acidovorax*), while others had negative effects (e.g. *Erwinia*). To the best of our knowledge, this is the first study explicitly describing the contribution of florally transmittable bacteria to seed communities and screening their effects on fruit set, seed germination and growth, which are important components of plant reproductive success.

### Floral transmission contributes to seed microbiota assembly, with potential roles for selective and stochastic processes

In our field survey, we found that on average 17% of the seed bacterial diversity was also found on stigmas, and that these shared ASVs make up 41% of the reads in seed communities on average (Fig. 2C). This is consistent with the range of overlap reported between flowers and seeds in *Phaseolus vulgaris* and *Raphanus sativus,* where 10% and 36% of seed bacterial taxa were shared, respectively (Chesneau *et al*., 2020). Our results build on this previous finding by showing that these shared bacteria make up over a third of the seed community in terms of abundance. While these contributions should be taken as estimates since we used different DNA extraction kits between tissue types (each with its own biases; Giangacomo *et al*., 2021), the large overlap is consistent with a contribution of floral transmission to bacterial community assembly in developing seeds. Furthermore, we validated floral transmission for four of the seven strains tested in our stigma inoculation experiment. Given the variable effects of the inoculated strains on fruit set, we propose that functionally diverse bacteria can colonize seeds via this pathway.

We also observed that bacterial richness was higher in stigmas compared to seeds (Fig. 1A) and that some shared genera were enriched in seeds compared to stigmas (Fig. 3B). Based on this, we suspect that there is a bottleneck during seed microbiota assembly (Newcombe *et al*., 2018). Since the shared taxa were distinct from those unique to stigmas (Fig. S4A) and only some of our bacterial strains colonized seeds from inoculated stigmas, this bottleneck could be due in part to selective filtering. This aligns with reports of maternal effects on seed microbiota assembly (Fort *et al*., 2021; Sulesky-Grieb *et al*., 2024). Such selection could be due to factors such as parent plant-bacterial interactions, resulting in transmission of bacteria with specific traits. This could help explain the transmission of *Erwinia* and *Rosenbergiella* (both belonging to the Enterobacterales, which are commonly associated with seeds; Simonin *et al*., 2022), as well as *Paraburkholderia* (previously shown to florally transmit to seeds; Mitter *et al*., 2017). It could also help explain why the seed-isolated *A. citrulli* and *Bacillus* strains were minimally or not detected respectively. It has been shown that *A. citrulli* can survive in seeds in a viable but nonculturable state (Kan *et al*., 2019), and some *Bacillus* species form spores that can be recalcitrant to culturing (Pious & Shaik, 2020). As such, it is possible that we were not able to sufficiently detect these strains using our culture-dependent approach.

We found that the overlap between seeds and stigmas from the same plant varied widely across plants and fields (Fig. 2B) and that there was high variability in bacterial re-isolation rates between seeds from different fruits and slices in the stigma inoculation experiment (Fig. 5A). These observations suggest a role for stochastic processes in floral transmission (Shade *et al*., 2017). This pattern is consistent with the hypothesis of imperfect bacterial transmission to seeds (Afkhami & Rudgers, 2008) and has been observed in culture-based studies of seed microbes (Newcombe *et al*., 2018; Shearin *et al*., 2018; Bergmann & Busby, 2021; Heitmann *et al*., 2021). Such stochastic variation could be due to several processes at different spatial scales. Given the ephemeral nature of flowers, the composition of the initial species pool on stigmas from different plants or fields would potentially dictate whether some taxa are successfully transmitted to seeds or not. Additionally, our observation of higher transmission rates in seeds closer to the stigma suggests dispersal limitation within fleshy fruits (i.e. at the micro-scale; Bergmann & Leveau, 2022). These findings open new opportunities for studying selective and stochastic mechanisms of seed microbiota assembly.

### Bee visitation did not affect seed bacterial community assembly

In our pollinator exclusion experiment, bee visitation did not introduce unique bacteria to seeds compared to pollen alone, suggesting pollination may vary in its effects on seed microbiota. This finding went against our expectations, as bee pollination has been previously shown to affect bacterial communities of flowers (Mcfrederick *et al*., 2017; Zemenick *et al*., 2018, 2021) and seeds (Prado *et al*., 2020). One of the most likely explanations for this is that we allowed bee visitation after hand pollinating stigmas, so any effect of bees would have to outweigh the effects of hand pollination. Additionally, we standardized the treatment to four bee visits per fruit, which is sufficient to fertilize seeds but much lower than potential visitation rates in the field (Stanghellini *et al*., 2002). However, commercial fields are more complex than a greenhouse or lab setting, so other factors (e.g. weather, handling by farm workers etc.) could impact the role of pollinators in vectoring bacteria. The standardization of bee visits in our experiment raises interesting questions about how visitation rates and visit quality could impact the incidence and extent of bacterial vectoring to seeds. We advocate for more experiments that demonstrate the extent of bees as a source of bacteria in the floral transmission pathway.

### Florally transmittable bacteria and their congeners impact fruit and seedling development

In our stigma inoculation experiment, we found that strains can either improve or reduce fruit set rates. For example, *Rosenbergiella sp.* P.s10.b1R significantly improved rates of successful fruit development (Fig. 5A). Additionally, *Microbacterium sp.* T.s9.b4R significantly improved fruit development rates while not being detected in seeds, suggesting that floral transmission is not required for stigma inoculation to affect fruit quality (i.e. legacy effects; Mallon *et al*., 2018). These positive effects could be due to bacterial production of plant hormones involved in fruit and seed development (e.g. gibberellins; Kumar *et al*., 2014; Bewley & Nonogaki, 2017; Eichmann *et al*., 2021). Alternatively, some bacteria could promote pollen germination (Christensen *et al*., 2021), thereby enhancing seed fertilization. The ability of bacteria to promote fruit set of *C. lanatus* is interesting as the basis for a greener alternative for chemicals that are used to promote fruit yield in melons (Hsu & Chen, 2024).

We also observed that seed inoculation with florally transmittable bacteria had variable effects on germination and seedling development. While *Rosenbergiella sp.* P.s10.b1R improved fruit development rates, it reduced seed germination and seedling emergence rates more than any other strain tested in our experiment. By contrast, *Rosenbergiella sp.* T.s1.b1 improved seedling development. We saw a similar pattern with our strains of *Paraburkholderia* and *Erwinia,* where one would improve germination and emergence while the other would reduce it. These patterns hint at functional variation of bacterial taxa between plant life stages and strains. We also observed that both *Acidovorax citrulli* strains had negligible or slightly positive effects on germination and emergence. Previous assays already showed that infected seedlings can be asymptomatic (Dutta *et al*., 2012b), and perhaps *A. citrulli* suppresses its pathogenic potential at this seedling stage to ensure dispersal at later plant life stages. This result supports the argument that the ecological role (e.g. pathogen, commensal, mutualist) of plant associated microbiota is context-dependent (Rodriguez *et al*., 2009; Leveau, 2024).

Overall, our findings here demonstrate that florally transmittable bacteria can have both positive and negative effects on seed and seedling fitness, depending on the strain. Combined with the fruit development data, these data indicate that floral transmission results in both beneficial and harmful bacteria colonizing seeds (Darsonval *et al*., 2008). Thus, our work highlights the importance of floral transmission both to seed bacterial community assembly and plant fitness.

**Table 4.** Bacterial genera isolated from stigmas and mature seeds of watermelon in 2021.

| Field | Tule |  | Poundstone |  | Total isolates (by genus) | Rel. abund. in stigmas (sequencing) | Rel. abund. in seeds (sequencing) | Isolates used in stigma experiment | Isolates used in seed experiment | Accessions |
| --- | --- | --- | --- | --- | --- | --- | --- | --- | --- | --- |
| *Genus* | Stigmas | Seeds | Stigmas | Seeds |  |  |  |  |  |  |
| *Bacillus* | 3 | 7* | 1 | 16 | 27 | 2.331 | 1.295 | T.m10.b1 | NA | PQ804665-PQ804691 |
| *Erwinia* | 1* | 0 | 7* | 0 | 8 | 0.079 | 0.779 | P.s8.b2 | P.s8.b2, P.s1.b3 | PQ804692-PQ804699 |
| *Rosenbergiella* | 1* | 0 | 7* | 0 | 8 | 1.375 | 0.511 | P.s10.b1 | P.s10.b1, T.s1.b1 | PQ804710-PQ804717 |
| *Paraburkholderia* | 3* | 0 | 0 | 0 | 3 | 0.122 | 0.021 | T.s8.b1 | T.s8.b1, T.s1.b2 | PQ804703-PQ804705 |
| *Priestia (formerly Bacillus)* | 1 | 1 | 0 | 1 | 3 | 2.331 | 1.295 | NA | NA | PQ804706-PQ804708 |
| *Mammaliicoccus (formerly Staphylococcus)* | 0 | 0 | 1 | 0 | 1 | 0.346 | 0 | NA | NA | PQ804700 |
| *Microbacterium* | 1* | 0 | 0 | 0 | 1 | 0.251 | 0.768 | T.s9.b4 | NA | PQ804701 |
| *Paenibacillus* | 0 | 0 | 1 | 0 | 1 | 0.221 | 0.029 | NA | NA | PQ804702 |
| *Ralstonia* | 0 | 1 | 0 | 0 | 1 | 9.638 | 0.041 | NA | NA | PQ804709 |
| *Sphingobium* | 0 | 0 | 0 | 1 | 1 | 0.02 | 0.457 | NA | NA | PQ804718 |
| **Total isolates (by source)** | **10** | **9** | **17** | **18** | **54** |  |  |  |  |  |

## Supporting information

Supplemental figures and tabbles

## Acknowledgements

We thank Dr. Amber Vinchesi-Vahl, Dr. Zheng Wang and Dr. Patricia Lazicki of the University of California Cooperative Extension service for coordinating communication with watermelon growers and providing access to the fields sampled in this study, along with field metadata. We thank Eduardo Ruiz for his assistance collecting samples from the field for the culture collection and the sequence-based survey. We thank Rudy Mahajan for his assistance in generating the culture collection. We thank Karen Barragan for her assistance on the pollinator exclusion experiment. We thank Dr. Ron Walcott, Dr. Bhabesh Dutta and Dr. Mei Zhao for providing *Acidovorax citrulli* strains for use in the inoculation experiments. Finally, we thank the Leveau Lab, the Vannette Lab and Dr. Marcel Holyoak for their feedback on manuscript drafts. We thank Muriel Bahut (ANAN platform, SFR Quasav) for *gyrB* amplicon sequencing.

## Data availability

Archives of strains from the field culture collection are available through the French Collection of Plant-Associated Bacteria (CIRM-CFBP). All sequences from the field culture collection were submitted to NCBI Genbank under accessions PQ804665-PQ804718. Raw sequences from the field survey and pollinator exclusion experiment were submitted to the NCBI Sequence Read Archive under BioProject PRJNA1208111. Metadata, datasets, R scripts and outputs are available at https://github.com/gbergmann/Bergmann.et.al_Floral_transmission_contribution_MS_2025.

## Funding

This work was supported by an NSF Graduate Research Fellowship awarded to GEB, a graduate research scholarship from the Sonoma County Mycological Association awarded to GEB and a UC Davis Jastro & Shields Graduate Award awarded to GEB. The seed viability experiment conducted by CL was supported by the Woodland Community College-MESA Undergraduate Research Program.

## Competing interests

None declared.

## Author contributions

GEB designed the study, led sampling and sample processing, DNA extractions, bioinformatic processing of sequence data and analysis, designed and conducted the stigma inoculation experiment, designed the seed viability assay, and wrote the manuscript. KL assisted on DNA extraction and sequence processing of the culture collection, and conducted part of the seed viability assay. CL conducted part of the seed viability assay. AV assisted in DNA extractions and sequence processing of the culture collection. CM prepared and sequenced the *gyrB* and 16S rRNA libraries of the pollinator experiment. MB and MS assisted in bioinformatic processing of the *gyrB* libraries, advised on data visualization and analysis, and revised the manuscript. RLV and JHJL provided guidance on study design, data analysis and figure presentation, and revised the manuscript.

## Descriptions of supplemental figures

**Table S1. Metadata collected during field sampling.**

**Figure S1. Faith’s phylogenetic diversity of stigma and seed communities**

**Figure S2. Contributions of shared (putative florally transmitted) taxa to richness (A) and reads (B) in stigma communities.**

**Figure S3. Nestedness vs. turnover in beta-diversity (A) and taxonomic composition (B) between stigma and seed communities.** The components of beta-diversity calculated in (A) were based off the prevalence-based Jaccard metric. Samples are grouped by field and individual plant in (B).

**Figure S4. Abundant taxa that are unique to stigmas (A) and seeds (B).**

**Figure S5. Microbiota composition varies highly in both hand-pollinated and insect-pollinated seeds.** (A) Relative abundances of the top 20 most abundant genera across pollination treatments. (B) beta-dispersion of seed communities between pollination treatments.

## Notes

https://github.com/gbergmann/Bergmann.et.al_Floral_transmission_contribution_MS_2025.

### Competing Interest Statement

The authors have declared no competing interest.

## References

Abdelfattah A, Tack AJM, Lobato C, Wassermann B, Berg G. 2022. From seed to seed: the role of microbial inheritance in the assembly of the plant microbiome. Trends in Microbiology 31: 346–355.

Abdelfattah A, Wisniewski M, Schena L, Tack AJM. 2021. Experimental evidence of microbial inheritance in plants and transmission routes from seed to phyllosphere and root. Environmental Microbiology 23: 2199–2214.

Afkhami ME, Rudgers JA. 2008. Symbiosis lost: Imperfect vertical transmission of fungal endophytes in grasses. The American Naturalist 172: 405–416.

Aizenberg-Gershtein Y, Izhaki I, Halpern M. 2013. Do honeybees shape the bacterial community composition in floral nectar? PLOS ONE 8: e67556.

Ambika Manirajan B, Ratering S, Rusch V, Schwiertz A, Geissler-Plaum R, Cardinale M, Schnell S. 2016. Bacterial microbiota associated with flower pollen is influenced by pollination type, and shows a high degree of diversity and species-specificity: The bacterial microbiota of flower pollen. Environmental Microbiology 18: 5161–5174.

Barret M, Briand M, Bonneau S, Préveaux A, Valière S, Bouchez O, Hunault G, Simoneau P, Jacques M-A. 2015. Emergence shapes the structure of the seed microbiota (HL Drake, Ed.). Applied and Environmental Microbiology 81: 1257–1266.

Baselga A, Orme CDL. 2012. betapart: an R package for the study of beta diversity. Methods in Ecology and Evolution 3: 808–812.

Beckers B, Op De Beeck M, Thijs S, Truyens S, Weyens N, Boerjan W, Vangronsveld J. 2016. Performance of 16s rDNA primer pairs in the study of rhizosphere and endosphere bacterial microbiomes in metabarcoding studies. Frontiers in Microbiology 7: 650.

Bergmann GE, Busby PE. 2021. The core seed mycobiome of *Pseudotsuga menziesii* var. *menziesii* across provenances of the Pacific Northwest, USA. Mycologia 113: 1169–1180.

Bergmann GE, Leveau JHJ. 2022. A metacommunity ecology approach to understanding microbial community assembly in developing plant seeds. Frontiers in Microbiology 13: 877519.

Bewley JD, Nonogaki H. 2017. Seed maturation and germination. Reference Module in Life Sciences: B9780128096338050925.

Bintarti AF, Sulesky-Grieb A, Stopnisek N, Shade A. 2022. Endophytic microbiome variation among single plant seeds. Phytobiomes Journal 6: 45–55.

Burdman S, Kots N, Kritzman G, Kopelowitz J. 2005. Molecular, physiological, and host-range characterization of *Acidovorax avenae* subsp. *citrulli* isolates from watermelon and melon in Israel. Plant Disease 89: 1339–1347.

Burdman S, Walcott RON. 2012. Acidovorax citrulli: generating basic and applied knowledge to tackle a global threat to the cucurbit industry. Molecular plant pathology 13: 805–815.

Busby PE, Newcombe G, Neat AS, Averill C. 2022. Facilitating reforestation through the plant microbiome: Perspectives from the phyllosphere. Annual Review of Phytopathology 60: 337– 356.

Busby PE, Soman C, Wagner MR, Friesen ML, Kremer J, Bennett A, Morsy M, Eisen JA, Leach JE, Dangl JL. 2017. Research priorities for harnessing plant microbiomes in sustainable agriculture. PLOS Biology 15: e2001793.

Callahan BJ, McMurdie PJ, Rosen MJ, Han AW, Johnson AJA, Holmes SP. 2016. DADA2: High-resolution sample inference from Illumina amplicon data. Nature Methods 13: 581–583.

Campbell JW, Daniels JC, Ellis JD. 2018. Fruit set and single visit stigma pollen deposition by managed bumble bees and wild bees in Citrullus lanatus (Cucurbitales: Cucurbitaceae). Journal of economic entomology 111: 989–992.

Chesneau G, Laroche B, Préveaux A, Marais C, Briand M, Marolleau B, Simonin M, Barret M. 2022. Single seed microbiota: Assembly and transmission from parent plant to seedling. mBio 13: e01648–22.

Chesneau G, Torres-Cortes G, Briand M, Darrasse A, Preveaux A, Marais C, Jacques M-A, Shade A, Barret M. 2020. Temporal dynamics of bacterial communities during seed development and maturation. FEMS Microbiology Ecology 96: fiaa190.

Christensen SM, Munkres I, Vannette RL. 2021. Nectar bacteria stimulate pollen germination and bursting to enhance microbial fitness. Current Biology 31: 4374–4380.

Conway JR, Lex A, Gehlenborg N. 2017. UpSetR: an R package for the visualization of intersecting sets and their properties (J Hancock, Ed.). Bioinformatics 33: 2938–2940.

Cui Z, Huntley RB, Schultes NP, Steven B, Zeng Q. 2020a. Inoculation of stigma-colonizing microbes to apple stigmas alters microbiome structure and reduces the occurrence of fire blight disease. Phytobiomes Journal 5: 156–165.

Cui Z, Huntley RB, Zeng Q, Steven B. 2020b. Temporal and spatial dynamics in the apple flower microbiome in the presence of the phytopathogen *Erwinia amylovora*. The ISME Journal 15: 318–329.

Darrasse A, Barret M, Cesbron S, Compant S, Jacques M-A. 2018. Niches and routes of transmission of *Xanthomonas citri* pv. *fuscans* to bean seeds. Plant and Soil 422: 115–128.

Darsonval A, Darrasse A, Meyer D, Demarty M, Durand K, Bureau C, Manceau C, Jacques M-A. 2008. The Type III secretion system of Xanthomonas fuscans subsp. fuscans is involved in the phyllosphere colonization process and in transmission to seeds of susceptible beans. Applied and Environmental Microbiology 74: 2669–2678.

Davis NM, Proctor DM, Holmes SP, Relman DA, Callahan BJ. 2018. Simple statistical identification and removal of contaminant sequences in marker-gene and metagenomics data. Microbiome 6: 226.

Debray R, Herbert RA, Jaffe AL, Crits-Christoph A, Power ME, Koskella B. 2022. Priority effects in microbiome assembly. Nature Reviews Microbiology 20: 109–121.

Du J, Liu Y, Zhu H. 2022. Genome-based analyses of the genus Acidovorax: proposal of the two novel genera Paracidovorax gen. nov., Paenacidovorax gen. nov. and the reclassification of Acidovorax antarcticus as Comamonas antarctica comb. nov. and emended description of the genus Acidovorax. Archives of Microbiology 205: 42.

Dutta B, Avci U, Hahn MG, Walcott RR. 2012a. Location of *Acidovorax citrulli* in infested watermelon seeds is influenced by the pathway of bacterial invasion. Phytopathology 102: 461– 468.

Dutta B, Gitaitis R, Sanders H, Booth C, Smith S, Langston DB. 2014. Role of blossom colonization in pepper Seed Infestation by *Xanthomonas euvesicatoria*. Phytopathology 104: 232–239.

Dutta B, Scherm H, Gitaitis RD, Walcott RR. 2012b. *Acidovorax citrulli* seed inoculum load affects seedling transmission and spread of bacterial fruit Bbotch of watermelon under greenhouse conditions. Plant Disease 96: 705–711.

Eichmann R, Richards L, Schäfer P. 2021. Hormones as go-betweens in plant microbiome assembly. The Plant Journal 105: 518–541.

Fort T, Pauvert C, Zanne A, Ovaskainen O, Caignard T, Barret M, Compant S, Hampe A, Delzon S, Vacher C. 2021. Maternal effects shape the seed mycobiome in *Quercus petraea*. New Phytologist 230: 1594–1608.

Free JB. 1993. Insect pollination of crops. London: Academic Press.

Giangacomo C, Mohseni M, Kovar L, Wallace JG. 2021. Comparing DNA Extraction and 16S rRNA Gene Amplification Methods for Plant-Associated Bacterial Communities. Phytobiomes Journal 5: 190–201.

Heitmann S, Bergmann GE, Barge E, Ridout M, Newcombe G, Busby PE. 2021. Culturable seed microbiota of *Populus trichocarpa*. Pathogens 10: 653.

Herrera CM, García IM, Pérez R. 2008. Invisible floral larcenies: Microbial communities degrade floral nectar of bumble bee-pollinated plants. Ecology 89: 2369–2376.

Heuer H, Krsek M, Baker P, Smalla K, Wellington EM. 1997. Analysis of actinomycete communities by specific amplification of genes encoding 16S rRNA and gel-electrophoretic separation in denaturing gradients. Applied and Environmental Microbiology 63: 3233–3241.

Hodgson S, de Cates C, Hodgson J, Morley NJ, Sutton BC, Gange AC. 2014. Vertical transmission of fungal endophytes is widespread in forbs. Ecology and Evolution 4: 1199–1208.

Hsu H-C, Chen W-L. 2024. CPPU Improves Fruit Setting and Growth in Greenhouse-grown Oriental Melons (Cucumis melo L. var. makuwa Makino). HortScience 59: 340–347.

Johnson KL, Minsavage GV, Le T, Jones JB, Walcott RR. 2011. Efficacy of a nonpathogenic Acidovorax citrulli strain as a biocontrol seed treatment for bacterial fruit blotch of cucurbits. Plant Disease 95: 697–704.

Johnson KL, Walcott RR. 2013. Quorum sensing contributes to seed-to-seedling transmission of Acidovorax citrulli on watermelon. Journal of Phytopathology 161: 562–573.

Johnston-Monje D, Gutiérrez JP, Lopez-Lavalle LAB. 2021. Seed-transmitted bacteria and fungi dominate juvenile plant microbiomes. Frontiers in Microbiology 12: 2945.

Kan Y, Jiang N, Xu X, Lyu Q, Gopalakrishnan V, Walcott R, Burdman S, Li J, Luo L. 2019. Induction and Resuscitation of the Viable but Non-culturable (VBNC) State in Acidovorax citrulli, the Causal Agent of Bacterial Fruit Blotch of Cucurbitaceous Crops. Frontiers in Microbiology 10: 1081.

Karstens L, Asquith M, Davin S, Fair D, Gregory WT, Wolfe AJ, Braun J, McWeeney S. 2019. Controlling for Contaminants in Low-Biomass 16S rRNA Gene Sequencing Experiments. mSystems 4: 10.1128/msystems.00290-19.

Khalaf EM, Raizada MN. 2016. Taxonomic and functional diversity of cultured seed associated microbes of the cucurbit family. BMC microbiology 16: 131.

Khalaf EM, Raizada MN. 2018. Bacterial seed endophytes of domesticated cucurbits antagonize fungal and oomycete pathogens including powdery mildew. Frontiers in microbiology 9: 42.

Kim H, Kim C, Lee Y-H. 2023. The single-seed microbiota reveals rare taxa-associated community robustness. Phytobiomes Journal 7: 324–338.

King EO, Ward MK, Raney DE. 1954. Two simple media for the demonstration of pyocyanin and fluorescin. The Journal of Laboratory and Clinical Medicine 44: 301–307.

Krassowski M. 2020. krassowski/complex-upset.

Kumar R, Khurana A, Sharma AK. 2014. Role of plant hormones and their interplay in development and ripening of fleshy fruits. Journal of Experimental Botany 65: 4561–4575.

Lahti L, Shetty S. 2017. Tools for microbiome analysis in R.

Lessl JT, Fessehaie A, Walcott RR. 2007. Colonization of female watermelon blossoms by *Acidovorax avenae ssp. citrulli* and the relationship between blossom inoculum dosage and seed infestation. Journal of Phytopathology 155: 114–121.

Leveau JHJ. 2024. Re-Envisioning the Plant Disease Triangle: Full Integration of the Host Microbiota and a Focal Pivot to Health Outcomes. Annual Review of Phytopathology 62: 31–47.

Lin H, Peddada SD. 2020. Analysis of compositions of microbiomes with bias correction. Nature Communications 11: 3514.

Lu W, Edelson J, Duthie J, Roberts B. 2003. A comparison of yield between high- and low-intensity management for three watermelon cultivars. HortScience 38.

Luo D-L, Huang S-Y, Ma C-Y, Zhang X-Y, Sun K, Zhang W, Dai C-C. 2024. Seed-borne bacterial synthetic community resists seed pathogenic fungi and promotes plant growth. Journal of Applied Microbiology 135: lxae073.

Mallon CA, Le Roux X, van Doorn GS, Dini-Andreote F, Poly F, Salles JF. 2018. The impact of failure: unsuccessful bacterial invasions steer the soil microbial community away from the invader’s niche. The ISME Journal 12: 728–741.

Mann LK. 1943. Fruit shape of watermelon as affected by placement of pollen on stigma. Botanical Gazette 105: 257–262.

Martin M. 2011. Cutadapt removes adapter sequences from high-throughput sequencing reads. EMBnet 17: 10–12.

Maude RB. 1996. Seedborne diseases and their control: principles and practice. Wallingford, Oxon, UK: CAB International.

Mcfrederick QS, Thomas JM, Neff JL, Vuong HQ, Russell KA, Hale AR, Mueller UG. 2017. Flowers and wild Megachilid bees share microbes. Microbial Ecology; Heidelberg 73: 188–200.

McMurdie PJ, Holmes S. 2013. phyloseq: An R Package for Reproducible Interactive Analysis and Graphics of Microbiome Census Data. PLOS ONE 8: e61217.

Mitter B, Pfaffenbichler N, Flavell R, Compant S, Antonielli L, Petric A, Berninger T, Naveed M, Sheibani-Tezerji R, von Maltzahn G, et al. 2017. A new approach to modify plant microbiomes and traits by introducing beneficial bacteria at flowering into progeny seeds. Frontiers in Microbiology 8: 11.

Murali A, Bhargava A, Wright ES. 2018. IDTAXA: a novel approach for accurate taxonomic classification of microbiome sequences. Microbiome 6: 140.

Murashige T, Skoog F. 1962. A Revised Medium for Rapid Growth and Bio Assays with Tobacco Tissue Cultures. Physiologia Plantarum 15: 473–497.

Newcombe G, Harding A, Ridout M, Busby PE. 2018. A hypothetical bottleneck in the plant microbiome. Frontiers in Microbiology 9: 1645.

Ngugi HK, Scherm H. 2004. Pollen mimicry during infection of blueberry flowers by conidia of *Monilinia vaccinii-corymbosi*. Physiological and Molecular Plant Pathology 64: 113–123.

Okansen J, Blanchet FG, Friendly M, Kindt R, Legendre P, McGlinn D, Minchin PR, O’Hara RB, Simpson GL, Solymos P, et al. 2019. vegan: Community ecology package.

Pal G, Kumar K, Verma A, Verma SK. 2022. Seed inhabiting bacterial endophytes of maize promote seedling establishment and provide protection against fungal disease. Microbiological Research 255: 126926.

Pious T, Shaik SP. 2020. Molecular Profiling on Surface-Disinfected Tomato Seeds Reveals High Diversity of Cultivation-Recalcitrant Endophytic Bacteria with Low Shares of Spore-Forming Firmicutes. Microbial ecology 79: 910–924.

Porter D. 1933. Watermelon breeding. Hilgardia 7: 585–624.

Prado A, Marolleau B, Vaissière BE, Barret M, Torres-Cortes G. 2020. Insect pollination: an ecological process involved in the assembly of the seed microbiota. Scientific Reports 10: 1–11.

Pruesse E, Quast C, Knittel K, Fuchs BM, Ludwig W, Peplies J, Glöckner FO. 2007. SILVA: a comprehensive online resource for quality checked and aligned ribosomal RNA sequence data compatible with ARB. Nucleic Acids Research 35: 7188–7196.

R Core Team. 2016. R: a language and environment for statistical computing.

Rebolleda-Gómez M, Ashman T-L. 2019. Floral organs act as environmental filters and interact with pollinators to structure the yellow monkeyflower (Mimulus guttatus) floral microbiome. Molecular Ecology 28: 5155–5171.

Rering CC, Beck JJ, Hall GW, McCartney MM, Vannette RL. 2018. Nectar-inhabiting microorganisms influence nectar volatile composition and attractiveness to a generalist pollinator. New Phytologist 220: 750–759.

Rodriguez RJ, White Jr JF, Arnold AE, Redman RS. 2009. Fungal endophytes: diversity and functional roles. New Phytologist 182: 314–330.

Schliep K, Paradis E, Martins L de O, Potts A, Bardel-Kahr I, White TW, Stachniss C, Kendall M, Halabi K, Bilderbeek R, et al. 2023. phangorn: Phylogenetic Reconstruction and Analysis.

Shade A, Jacques M-A, Barret M. 2017. Ecological patterns of seed microbiome diversity, transmission, and assembly. Current Opinion in Microbiology 37: 15–22.

Shearin ZRC, Filipek M, Desai R, Bickford WA, Kowalski KP, Clay K. 2018. Fungal endophytes from seeds of invasive, non-native Phragmites australis and their potential role in germination and seedling growth. Plant and Soil 422: 183–194.

Simonin M, Briand M, Chesneau G, Rochefort A, Marais C, Sarniguet A, Barret M. 2022. Seed microbiota revealed by a large-scale meta-analysis including 50 plant species. New Phytologist 234: 1448–1463.

Singh P, Vaishnav A, Liu H, Xiong C, Singh HB, Singh BK. 2023. Seed biopriming for sustainable agriculture and ecosystem restoration. Microbial Biotechnology 00: 1–11.

Stanghellini MS, Ambrose JT, Schultheis JR. 2002. Diurnal activity, floral visitation and pollen deposition by honey bees and bumble bees on field-grown cucumber and watermelon. Journal of Apicultural Research 41: 27–34.

Sulesky-Grieb A, Simonin M, Bintarti AF, Marolleau B, Barret M, Shade A. 2024. Stable, multigenerational transmission of the bean seed microbiome despite abiotic stress. mSystems 9: e0095124.

Tang X, Liu Q, Luo L, Yin C. 2023. The endophyte Bacillus amyloliquefaciens from Picea asperata seeds promotes seed germination and its physiological mechanism. Journal of Soil Science and Plant Nutrition 24: 421–434.

USDA/NASS. 2023. 2023 state agriculture overview for California.

Vannette RL, Gauthier M-PL, Fukami T. 2012. Nectar bacteria, but not yeast, weaken a plant-pollinator mutualism. Proceedings of the Royal Society B: Biological Sciences 280: 20122601– 20122601.

Walcott RR, Gitaitis RD, Castro AC. 2003. Role of blossoms in watermelon seed infestation by *Acidovorax avenae* subsp. *citrulli*. Phytopathology 93: 528–534.

Walcott RR, Langston DB, Sanders FH, Gitaitis RD. 2000. Investigating intraspecific variation of *Acidovorax avenae* subsp. *citrulli* using DNA fingerprinting and whole cell fatty acid analysis. Phytopathology® 90: 191–196.

Wijesinghe SAEC, Evans LJ, Kirkland L, Rader R. 2020. A global review of watermelon pollination biology and ecology: The increasing importance of seedless cultivars. Scientia Horticulturae 271: 109493.

Yao Y, Liu C, Zhang Y, Lin Y, Chen T, Xie J, Chang H, Fu Y, Cheng J, Li B, et al. 2024. The dynamic changes of Brassica napus seed microbiota across the entire seed life in the field. Plants 13: 912.

Zemenick AT, Rosenheim JA, Vannette RL. 2018. Legitimate visitors and nectar robbers of Aquilegia formosa have different effects on nectar bacterial communities. Ecosphere 9: e02459.

Zemenick AT, Vannette RL, Rosenheim JA. 2021. Linked networks reveal dual roles of insect dispersal and species sorting for bacterial communities in flowers. Oikos 130: 847376.

