## Supplemental figures and tabbles for "Contribution of floral transmission to the assembly and health impact of bacterial communities in watermelon seeds"

### Supplemental figures/tables

Table S1. Metadata of fields sampled sequence-based survey

| Field ID | Year sampled | Field county | Cultivar | # stigmas collected | # fruits collected | Stigma sampling date | Fruit sampling date | # of beehives |
|---|---|---|---|---|---|---|---|---|
| Co1 | 2022 | Colusa | Crimson Sweet | 10 | 10 | 7/15/2022 | 7/22/2022 | 1 |
| Co2 | 2022 | Colusa | AU producer | 10 | 10 | 8/4/2022-8/12/2022 | 8/12/2022-8/19/2022 | 8 |
| SJ1 | 2022 | San Joaquin | Sentinel | 10 | 10 | 6/24/2022 | 7/1/2022 | 4 |
| SJ2 | 2022 | San Joaquin | Sentinel | 10 | 10 | 7/7/2022 | 7/14/2022 | 1 |

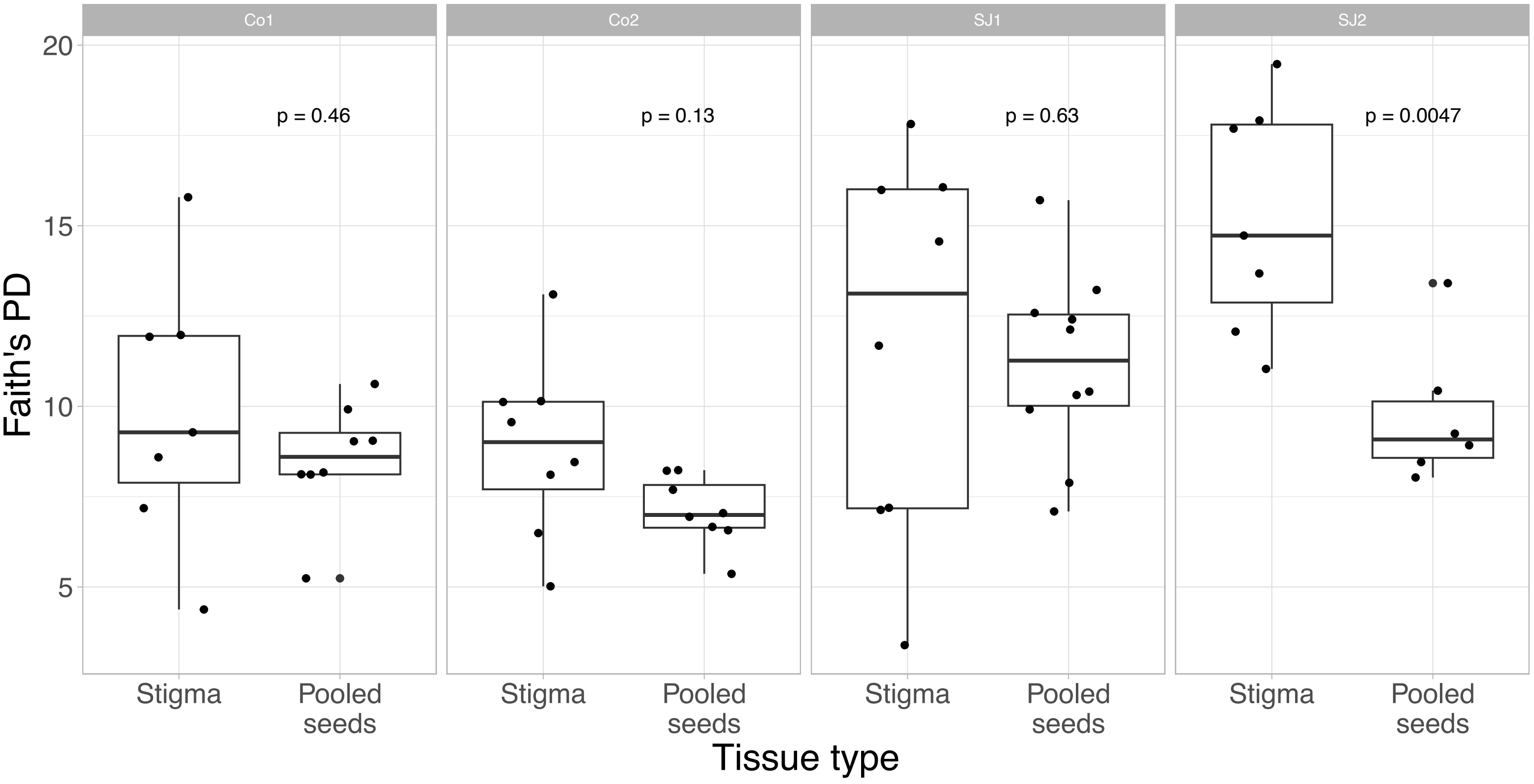


### Figure S1. Faith’s phylogenetic diversity of stigma and seed communities

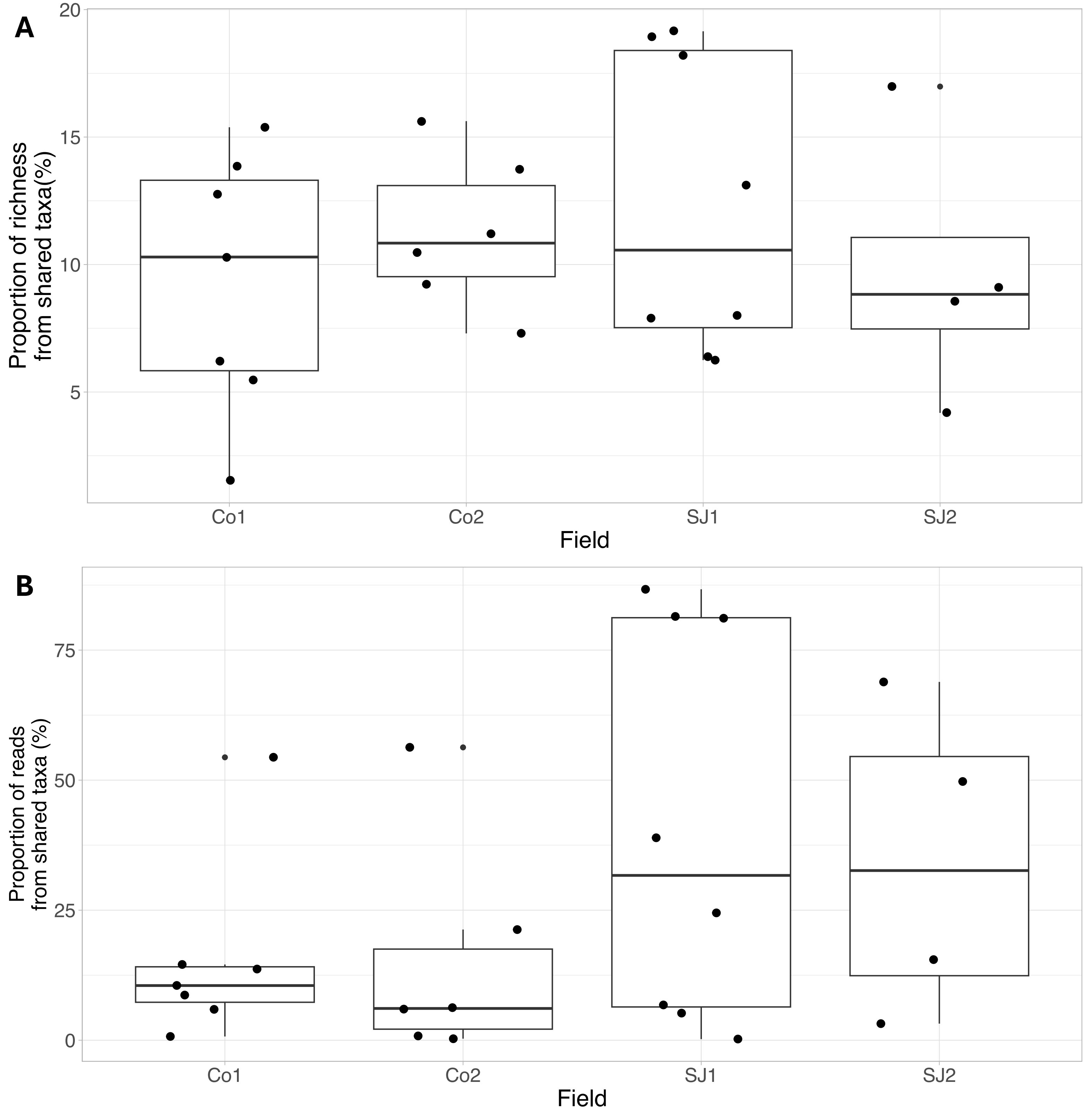


Figure S2: Contributions of shared taxa to richness (A) and reads (B) in stigma communities.

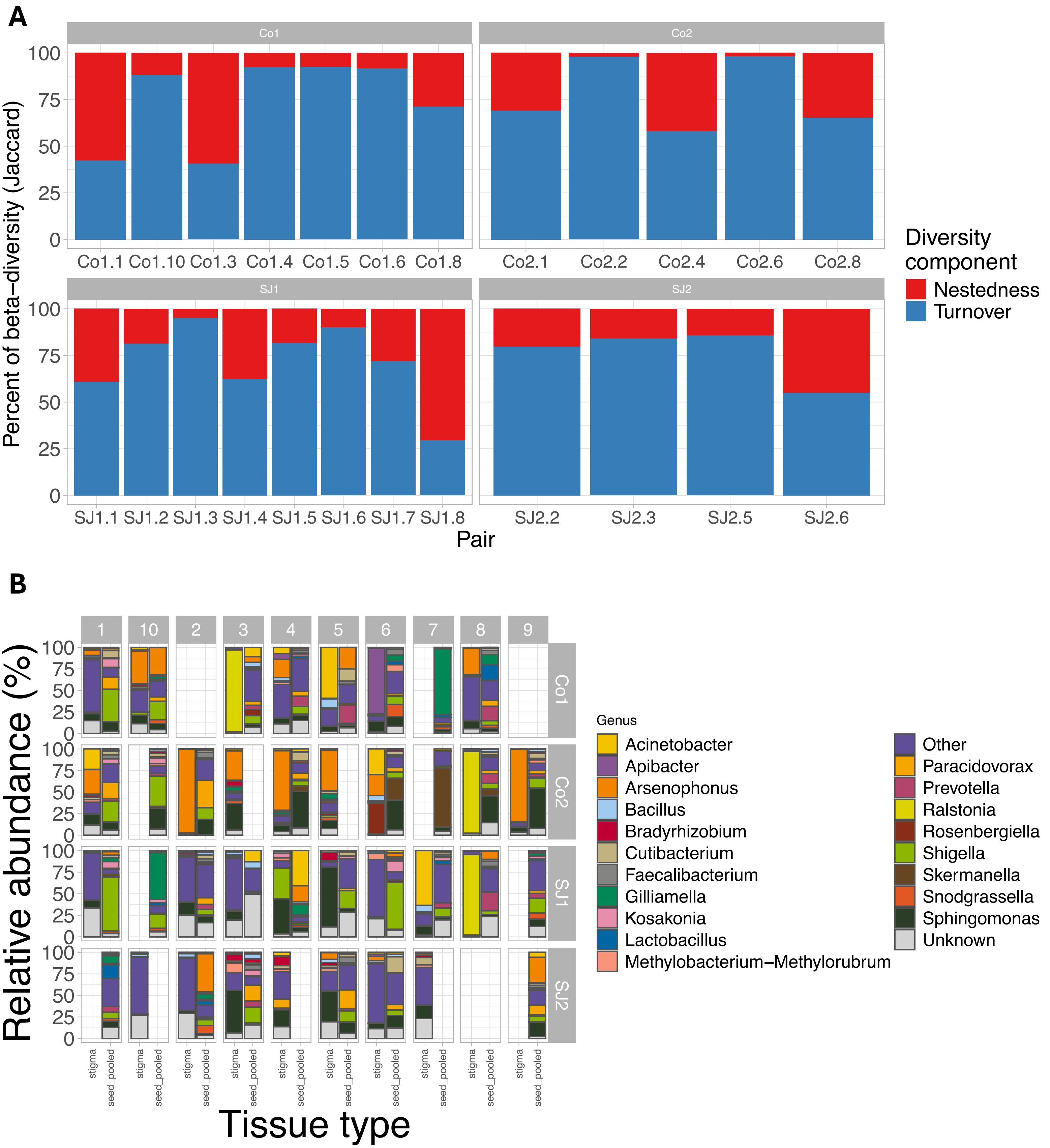


Figure S3. Nestedness vs. turnover in beta-diversity (A) and taxonomic composition (B) between stigma and seed communities. The components of beta-diversity calculated in (A) were based off the prevalence-based Jaccard metric. Samples are grouped by field and individual plant in (B).

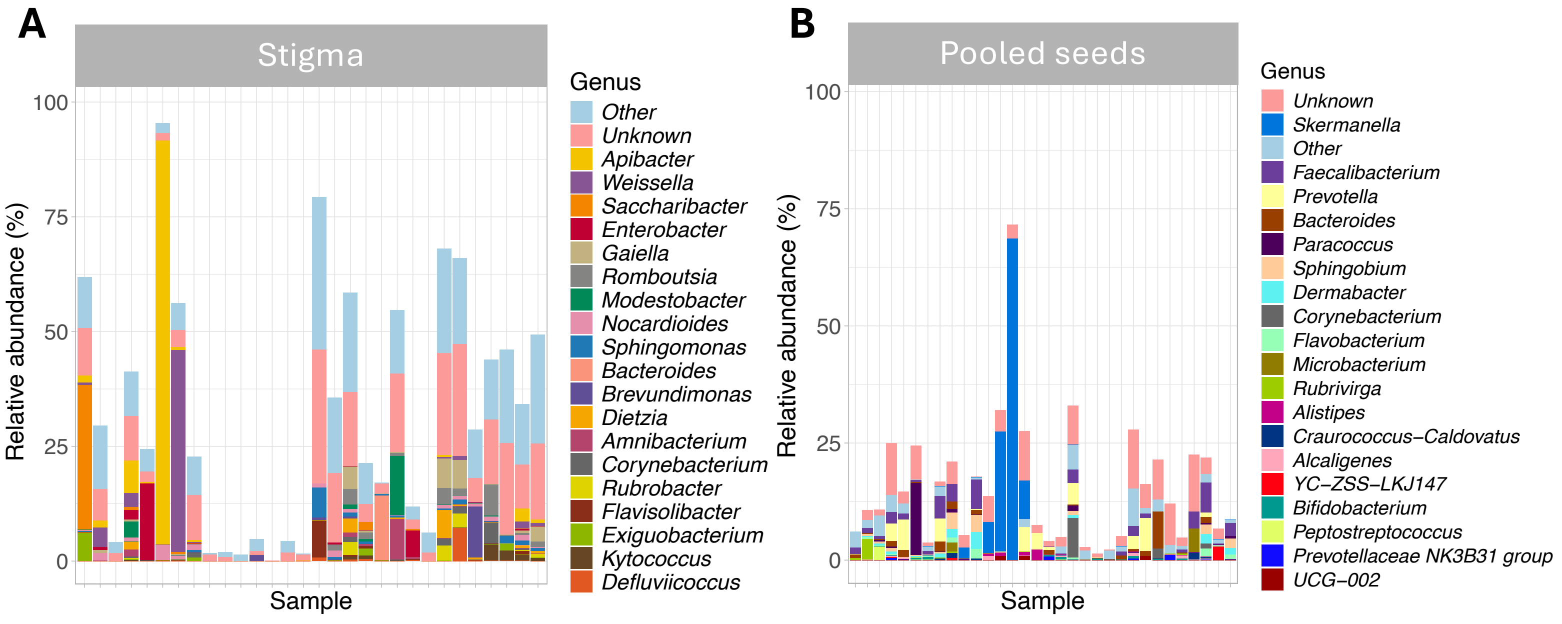


Figure S4: Abundant taxa that are unique to stigmas (A) and seeds (B)

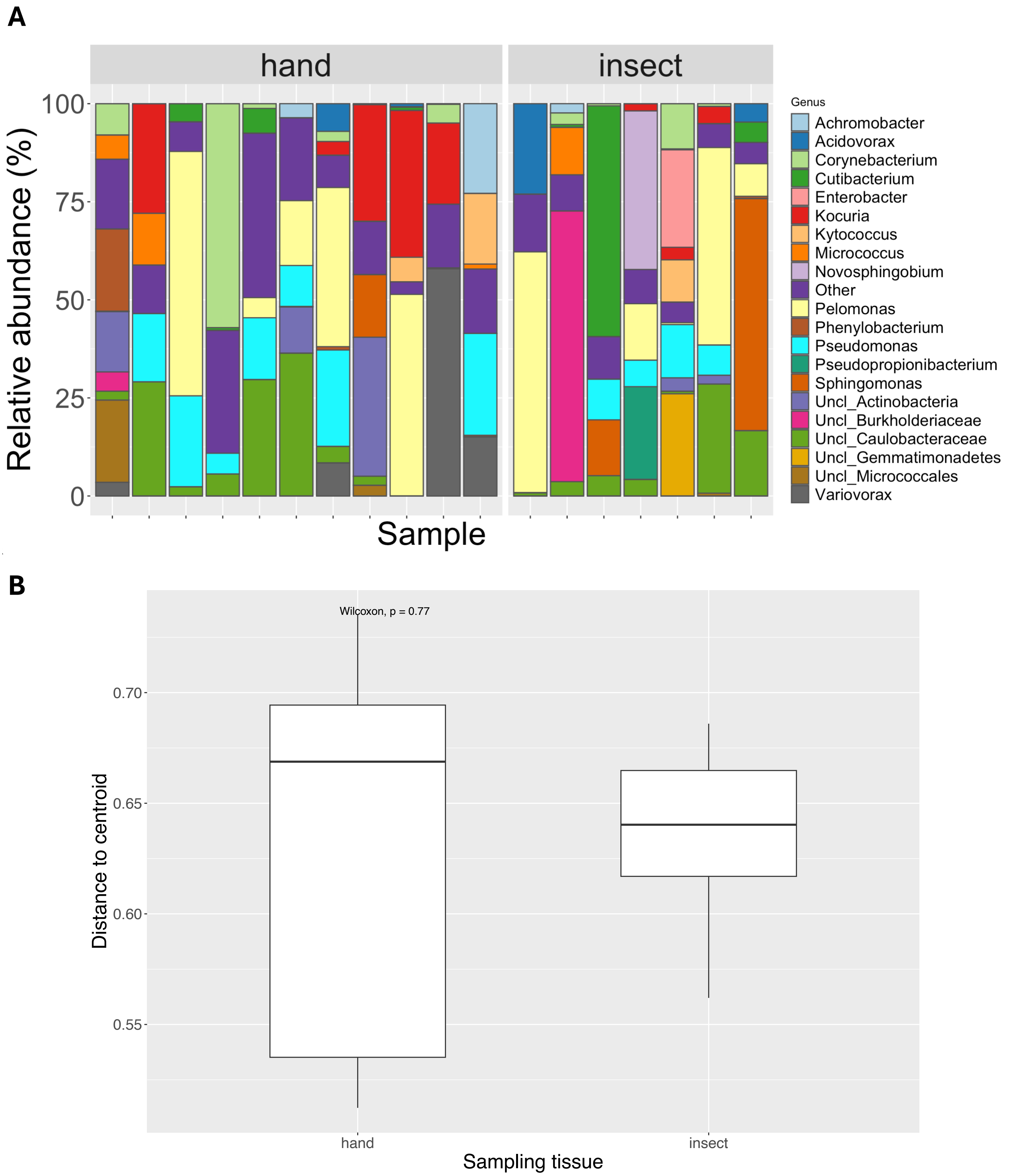


Figure S5: Microbiota composition varies highly in both hand-pollinated and insect-pollinated seeds. (A) Relative abundances of the top 20 most abundant genera across pollination treatments. (B) beta-dispersion of seed communities between pollination treatments.
